# Using Mathematical Modeling to Distinguish Intrinsic and Acquired Targeted Therapeutic Resistance in Head and Neck Cancer

**DOI:** 10.1101/2022.02.18.481078

**Authors:** Santiago D. Cardenas, Constance J. Reznik, Ruchira Ranaweera, Feifei Song, Christine H. Chung, Elana J. Fertig, Jana L. Gevertz

## Abstract

The promise of precision medicine has been limited by the pervasive therapeutic resistance to many targeted therapies for cancer. Inferring the timing (i.e., pre-existing or acquired) and mechanism (i.e., drug-induced) of such resistance is crucial for designing effective new therapeutics. This paper studies the mechanism and timing of cetuximab resistance in head and neck squamous cell carcinoma (HNSCC) using tumor volume data obtained from patient-derived tumor xenografts. We propose a family of mathematical models, with each member of the family assuming a different timing and mechanism of resistance. We present a method for fitting these models to individual volumetric data, and utilize model selection and parameter sensitivity analyses to ask: which member of the family of models best describes HNSCC response to cetuximab, and what does that tell us about the timing and mechanisms driving resistance? We find that along with time-course volumetric data to a single dose of cetuximab, the initial resistance fraction and, in some instances, dose escalation volumetric data are required to distinguish among the family of models and thereby infer the mechanisms of resistance. These findings can inform future experimental design so that we can best leverage the synergy of wet laboratory experimentation and mathematical modeling in the study of novel targeted cancer therapeutics.

## 1 Introduction

In cancer, each individual’s tumor has undergone a distinct set of molecular and cellular alterations that promote malignancy. Advances to high-throughput measurement technologies have enabled unprecedented characterization of these alterations, ushering in a new era of precision medicine which selects therapies to target the specific changes in each tumor. In spite of the promise of these precision medicine strategies, many cancers do not respond as anticipated to such targeted therapeutic strategies, and those who do respond frequently develop resistance.

Head and neck squamous cell carcinoma (HNSCC) is the 6th most common cancer world-wide with a 5-year survival rate of 50% [18]. Increased expression of the epidermal growth factor receptor (EGFR) occurs in 90% of HNSCC and is associated with poor survival [12, 39]. EGFR is a receptor in certain types of cells that binds to epidermal growth factors, which are involved in cell signaling pathways controlling cell division and survival. Therefore, targeted therapeutics inhibiting EGFR have been developed to block these pathways as a precision therapeutic to prevent cancer cells from growing. Cetuximab is the only targeted therapy FDA approved for HNSCC [45]. Including cetuximab as part of an advanced stage HNSCC treatment plan exhibits a survival advantage for the patient compared to radiation treatment alone. Cetuximab also improves response rates compared to chemotherapy in patients with metastatic or recurrent HNSCC [37]. However, only a subset of patients are intrinsically sensitive to cetuximab, and responsive patients will develop resistance within one to two years [45, 5, 53, 34]. The widespread prevalence of cetuximab resistance is currently limiting its clinical utility in HNSCC.

There are three different types of drug-resistance to consider in understanding resistance to targeted therapies such as cetuximab: pre-existing resistance, randomly-acquired resistance, and drug-induced acquired resistance. Pre-existing resistance is when all resistance in the tumor population is assumed to exist before treatment begins. Treatment then selects for these resistant cells, giving rise to a resistant tumor. Random acquired resistance occurs when resistant cells arise during treatment due to random genetic mutations or phenotypic switching, but not as a result of the drug administered. Cells for which resistance is pre-existing or randomly acquired act as a substrate for Darwinian evolution [3]. Lastly, drug-induced resistance is resistance directly caused by the drug during treatment, either through genetic changes or more likely through non-genetic cell phenotype plasticity [48, 43, 28, 3]. These cells, often called drug-resistant or drug-tolerant persisters, act as a substrate for Lamarckian evolution as the adaptive changes occur as a direct response to the drug itself [3]. Single-cell data of HNSCC cell lines suggests that the molecular mechanisms of compensatory growth factor signaling and epithelial to mesenchymal transition that underlie therapeutic resistance to cetuximab can be induced as an early response to treatment [29]. However, their precise contribution to subsequent resistance requires further longitudinal profiling which can be confounded by evolutionary processes in culture [50] and infeasible to extend to powered, temporal profiling in *in vivo* models.

Mathematical models have been widely utilized to help understand drug resistance, and its consequences for treatment response and design - see [51, 7, 17, 33] for reviews of modeling work on cancer drug resistance. Overwhelmingly, these models have assumed that resistance is either pre-existing (as in [26, 16, 49, 40]), or is a combination of pre-existing and spontaneously acquired resistance (as in [15, 30, 22]). More recently, modeling has also considered the contribution of the drug itself in driving the formation of resistance. Works such as [43, 21, 9, 2, 14] consider drug-induced resistance, though they are limited in their ability to make predictions regarding doses and dosages that differ from the data used to validate them, as these models are dose-independent. A handful of mathematical models have been developed in which resistance is induced by the drug itself in a dose-dependent fashion [10, 20, 35, 23]. The modeling family herein is strongly motivated by the single model proposed in [23], wherein pre-existing, spontaneously acquired, and dose-dependent drug-induced resistance are modeled through a minimal system of two ordinary differential equations.

In this work, we propose a family of mathematical models, with each “member” of the family assuming a different timing and mechanism of cetuximab resistance. In Section 2 we detail the protocol for collecting the experimental data, describe the family of mathematical models (where each member of the family represents a different set of mechanisms driving resistance), explain the algorithm for fitting these models to individual volumetric data, and introduce the methodology for assessing parameter sensitivity/identifiability. In Section 3 we employ information criteria (IC) to try and identify the “best” model to describe the data (that is, the model with the lowest IC value across the individual samples). This information theoretic approach determined that control growth can be well-described using a simple exponential model. Extending such an information theoretic approach to our family of resistance models allowed us to confidently conclude that the data cannot be explained without resistance, and that the combination of pre-existing and randomly acquired resistance is very unlikely to be mechanism responsible for the resistance to cetuximab observed in the experiments. In Section 3 we use a profile likelihood analysis to demonstrate that single-cell experiments which measure the resistance fraction in the initial tumor population provide powerful data for selecting the model (and therefore the underlying mechanisms) most parsimonious with the experimental data. In the case where this measure of pre-existing resistance does not allow the mechanism of resistance to be definitively determined using our family of models, we further propose that a dose-escalation experiment would provide the needed data to identify the model whose mechanisms best-explain the resistance observed to cetuximab. Section 4 contains closing remarks and reflections about the role mathematical modeling can play in experimental design to decipher the mechanism of resistance to targeted therapeutics.

## 2 Methods and Model

### 2.1 Experimental Data

In this project, we utilize tumor volume data obtained from temporally monitoring a cetuximab responsive patient-derived tumor xenograft HNSCC model. Tumor tissue were collected from surgically resected HNSCC patients under the auspices of a tissue bank protocol approved by Johns Hopkins University Institutional Review Board. All animal studies and care were approved by the Institutional Animal Care and Use Committee of the Johns Hopkins University and Moffitt Cancer Center. Following HNSCC tumor resection, de-identified patient samples were implanted into athymic nude mice (Crl: NU-Foxn1nu, 4–6 weeks old; 20 g; Harlan Laboratories, Indianapolis, IN) and passaged to subsequent generations of mice for expansion. The mice are then divided into two groups: the control group and the treatment group. For each group, tumor volume is tracked over time under the assumption that 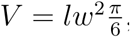, where *l* is length and *w* is width of the tumor. Treatment (either with a placebo, or with cetuximab) starts when the tumor volume is ~200 mm^3^. Mice were euthanized if tumor volume surpassed ~2,000 mm^3^, if they lost more than 25% of their body weight, or if ulceration occurred on the skin over the tumors.

The control data is obtained by administering a weekly dose of phosphate-buffered saline (PBS) to tumor-bearing mice. We classified each control mouse into one of three categories: increasing volume, decreasing volume, and stabilized volume. Out of 25 control mice, 19 show increasing volume (see Mouse 23 in Fig. 2), one shows decreasing volume (see Mouse 11 in Fig. 2), and five show stabilization (see Mouse 22 in Fig. 2).

The treatment data is obtained by following the same procedure as the control mice, except that mice were given a 5 mg/kg intraperitoneal injection of cetuximab once every 7 days. As with the control mice, each mouse was classified as either increasing in volume (treatment failure), decreasing in volume (treatment success), or stabilized volume. Out of 29 mice, 19 show increasing volume (see Mouse 13 in Fig. 4), seven show decreasing volume (see Mouse 23 in Fig. 4), and three show stabilization (see Mouse 24 in Fig. 4). A Fisher’s exact test has a p-value of 0.06 of decreased tumor volume occurring in the treatment group relative to control group, suggesting a trend towards Cetuximab response in this xenograft model.

To account for outliers and noise in our data, we applied a censor to remove any data points deemed not biologically plausible. Estimates from literature of doubling time for HNSCC vary widely, from 26 hours in culture to 44 days in vivo [19, 25]. According to exponential fits, in the control data the fastest cell-doubling time observed was 13 days. We took a conservative approach to censoring the data: in any case where the data show the tumor more than doubling in volume in 3-4 days, and the subsequent time points are not consistent with that rapid doubling time (meaning, the larger increase in volume is not sustained beyond that one point), we remove the outlier volume. We show two examples to depict our censoring approach in Fig. S1, one with a censored point due to an unsustained rapid doubling, and one without censoring despite a rapid doubling as it was sustained beyond that time point. In the control data, exactly one data point was removed from five of 25 mice, and in the treatment data nine data points were censored across seven of 29 mice.

### 2.2 Modeling Control Data

Before building a model of tumor growth in response to treatment, we first considered how to best-describe tumor growth in the absence of treatment. There are a multitude of mathematical equations to describe tumor growth, and the equation chosen can have important consequences on model predictions [46, 41]. Herein, we considered three different models of tumor growth: exponential, logistic, and Allee. These models were chosen because they represent a hierarchy of complexity.

Exponential growth simply assumes the growth rate of the tumor volume *V* is proportional to the tumor volume:

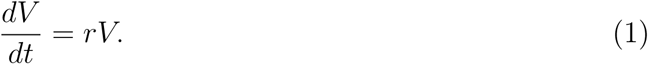

Logistic growth adds a rate-limiting factor to uncontrolled exponential growth, accounting for environmental constraints on tumor progression through a carrying capacity *K*:

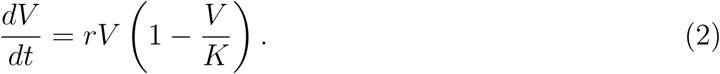

Finally, the Allee effect further adds the assumption that the growth rate can also be limited by a population size that is below the Allee threshold *m*:

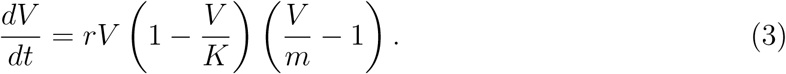

There are multiple plausible explanations for why tumor growth could be described by such a differential equation. It may be that tumors grow slower because of uptake challenges in the xenograft system, or because they have yet to accumulate significant mutations. As an example, the growth kinetics of BT-474 luminal B breast cancer cells was shown to be best-described by a model structure that considers the Allee effect [27].

### 2.3 Modeling Treatment Data

Once the “best” control model is selected, we can move to build a model that incorporates treatment response to cetuximab. We will use the following general modeling framework, where *S* is the volume of cells that are sensitive to cetuximab, *R* is the volume of cells with some level of resistance to cetuximab, and *D* is the concentration of drug:

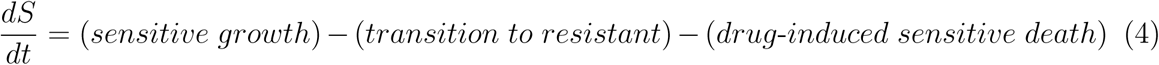

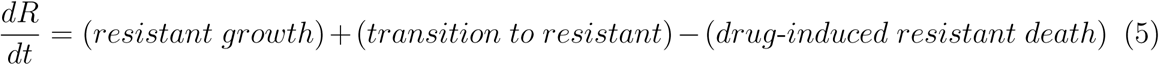

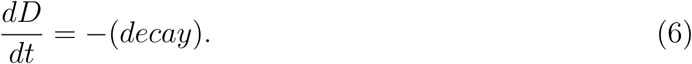

We assume that growth is exponential (justified in Section 3.1), that the death rate is proportional to the drug concentration and the volume of the subpopulation, and that in eqn. (6) the only dynamics modeled are the natural decay of the drug. This simplifies our general modeling framework to have the form:

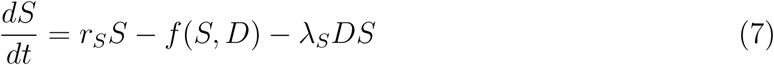

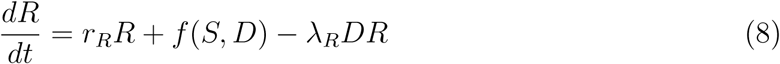

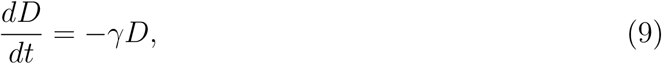

where *r*_*S*_ is the growth rate for sensitive cells, and *r*_*R*_ is the growth rate for resistant cells. We assume that *r*_*S*_ ≥ *r*_*R*_, as resistance will either not impact the growth rate of cells, or it will result in a fitness disadvantage [47, 6]. *λ*_*S*_ is the drug-induced death term for sensitive cells, and *λ*_*R*_ is the drug-induced death term for resistant cells. We assume that *λ*_*S*_ *> λ*_*R*_, as by definition, sensitive cells must be easier for the drug to kill. *f* (*S, D*) is the function that represents the transition of sensitive cells to resistant ones (i.e., acquired resistance). This may or may not depend on the drug *D*. Finally, *γ* is decay rate of the drug, which we fix using the the fact that the mean half-life of cetuximab is 4.75 days [1], corresponding to *γ* ≈ 0.1459 days^−1^.

Depending on the assumptions made, this model can represent any combination of: pre-existing resistance (when *R*(0) *>* 0), randomly acquired resistance (when the transition to resistance *f* (*S, D*) is independent of the drug *D*), and drug-induced acquired resistance (when *f* (*S, D*) depends on *D*). The different sub-models that we consider are explained here, and visually explained in Fig. 1.

**Figure 1:**
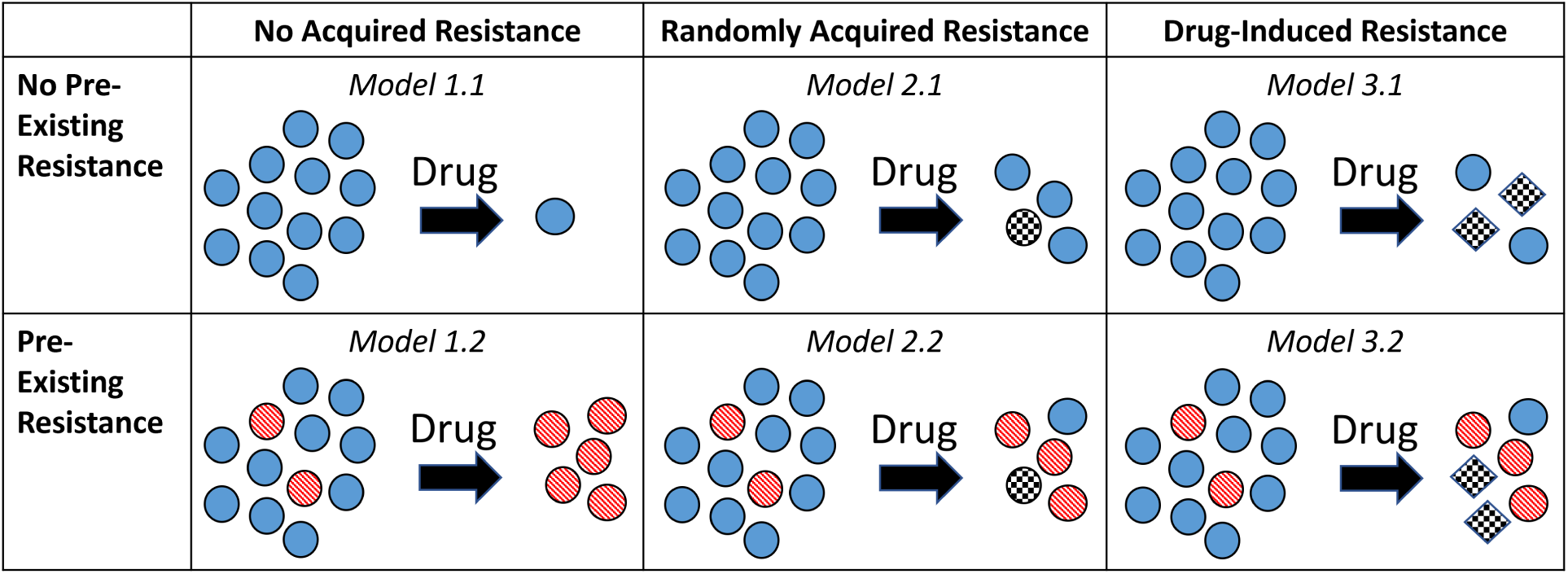
Schematic illustrating the family of resistance models. Sensitive cells are illustrated as blue circles, pre-existing resistant cells as red-striped circles, spontaneously-created resistant cells as black-and-white checkered circles, and drug-induced resistant cells as black-and-white checkered diamonds. Second row contains all models without pre-existing resistance, and the third row contains all models with pre-existing resistance. Second column contains all models with no acquired resistance, third column contains all models with randomly acquired resistance, and fourth column contains all models with drug-induced resistance.

- **Model 1: No Acquired Resistance**. This requires setting *f* (*S, D*) = 0 in eqns. (7)-(8). This model can be further broken down into two sub-cases:
  – **Model 1.1: No Pre-Existing Resistance**. Achieved by setting *R*(0) = 0, meaning the entire tumor population is sensitive to cetuximab.
  – **Model 1.2: Pre-Existing Resistance**. Achieved by allowing *R*(0) *>* 0.
- **Model 2: Randomly Acquired Resistance**. We model this with a random transition term *f* (*S, D*) = *f* (*S*) = *gS* in eqns. (7)-(8). This model can be further broken down into two sub-cases:
  – **Model 2.1: No Pre-Existing Resistance**. Achieved by setting *R*(0) = 0, meaning resistance can only result from the random acquisition of resistance that happen during treatment.
  – **Model 2.2: Pre-Existing Resistance**. Achieved by allowing *R*(0) *>* 0, meaning resistance can pre-exist treatment and can be randomly acquired during treatment.
- **Model 3: Drug-Induced Acquired Resistance**. We model this with a drug-dependent transition term *f* (*S, D*) = *gSD* in eqns. (7)-(8), as similarly done in [23]. This model can be further broken down into two sub-cases:
  – **Model 3.1: No Pre-Existing Resistance**. Achieved by setting *R*(0) = 0, meaning resistance can only result from the drug-induced acquisition of resistance.
  – **Model 3.2: Pre-Existing Resistance**. Achieved by allowing *R*(0) *>* 0, meaning resistance can pre-exist treatment and can be induced by the drug during treatment.

### 2.4 Fitting Algorithm

As we have proposed a variety of models to describe resistance of HNSCC to cetuximab, we must determine which model (or models) most accurately describes the experimental data. That is, we must fit each model to the volumetric time-course data of treatment response to cetuximab. Due to the extreme variability between mice, we chose to fit each mouse individually, rather than fit to the average of the time-course data.

For each model, and for each mouse *i*, the parameter set we seek is the one that minimizes the sum of the squared error (SSE), which we will call *ζ*_*i*_:

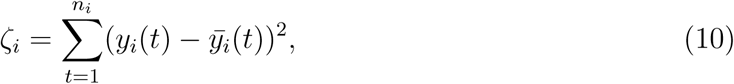

where *y*_*i*_(*t*) represents the experimental tumor volume for mouse *i* at time *t*, and 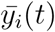 is the tumor volume obtained through the model at time *t*. This is indexed over *n*_*i*_, the number of time points in the data set for mouse *i*.

We implement a two-step fitting approach. The first step uses a Quasi-Monte Carlo (QMC) method to randomly sample the parameter space. Herein we use Sobol’s low-discrepancy sequences to uniformly place the randomly-sampled points across a *k*-dimensional hyperrectangle [31]. *k*-dimensions represents the *k* parameters in the model that are being fit to the data (including the initial condition *S*(0), and *R*(0) in the case of pre-existing resistance).

Our algorithm utilizes QMC by first randomly sampling 1.5 *×* 10^6^ Sobol points of the form (*p*_1_, …, *p*_*k*_). We chose this number of Sobol points to minimize the computational time required while maximizing coverage of the parameter space. Each *p*_*i*_ in a sampled point are in the range [0, 1]. We then have to scale the values of *p*_*i*_ so that they are in a biologically reasonable range for that parameter value. Somewhat arbitrarily, we found it sufficient to scale all non-initial condition and non-carrying capacity parameters except for *r*_*S*_ to be in the range [0, 0.1], as parameter values beyond this result in model predictions of completely different magnitudes than the experimental data. Numerical experimentation suggested optimal *r*_*S*_ values to be greater than 0.1, so this parameter was scaled to be in the range [0, 0.2]. The scaled range for the initial tumor volume was mouse-dependent. The initial condition per mouse were scaled to be in the range [0, 2*V*_0_], where *V*_0_ is the actual initial tumor volume for each mouse (see Table S1 in the Appendix). Further, the carrying capacity was searched over the range [*V*_0_, 10^5^]. Finally, in the case of the Allee effect, the existential threshold *m* was searched over the range [0, 10*V*_0_].

The model of interest is then solved at the 1.5 × 10^6^ scaled Sobol parameter sets for a given mouse *i*, provided the parameter set is biologically realistic. Biological viability is determined by the restrictions detailed in Section 2.3 based on the fitness disadvantage conferred by drug resistance (*r*_*S*_ ≥ *r*_*R*_ and *λ*_*S*_ *> λ*_*R*_). Each biologically viable parameter set provides us with a cost function value, *ζ*_*i*_(*p*_1_, …, *p*_*k*_). Then, for each mouse *i* we identify the parameter set with the lowest *ζ*_*i*_ value. This parameter set should be close to the optimal parameter set, though generally it is not the actual optimal. The Quasi-Monte Carlo step is summarized in Algorithm 1.

#### Algorithm 1: Quasi-Monte Carlo Method

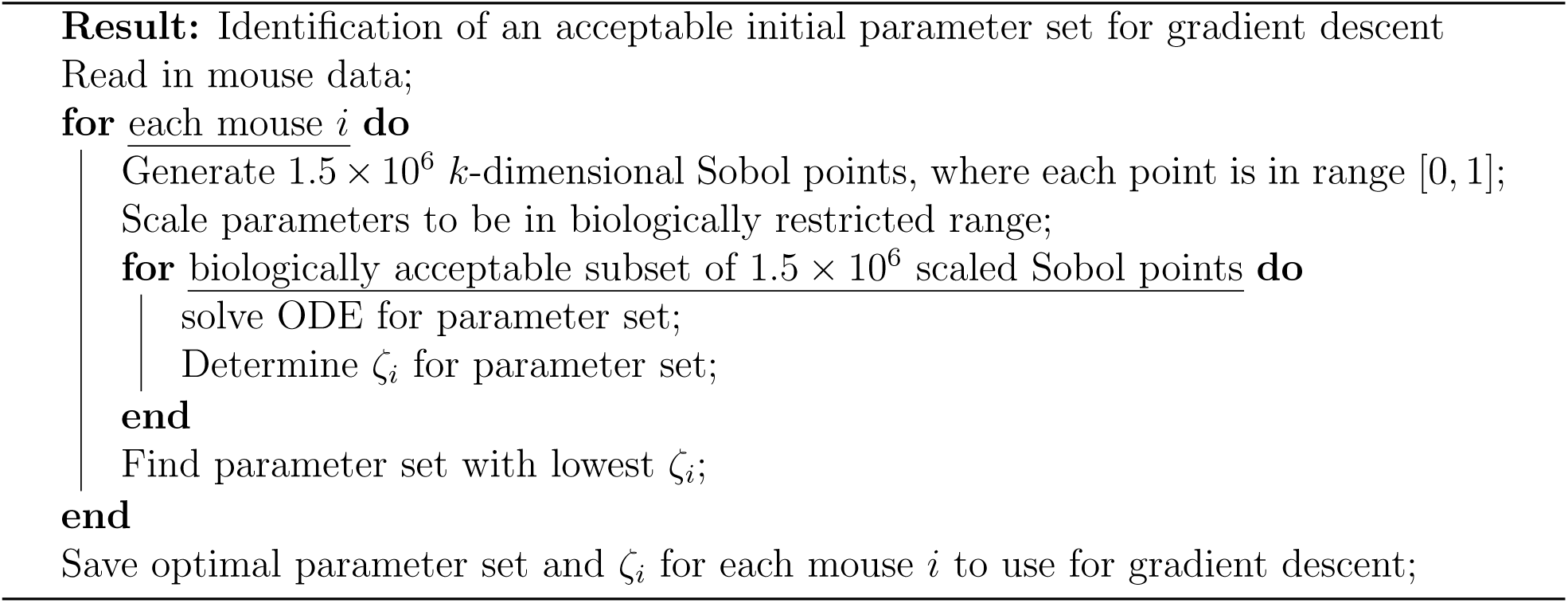

Once QMC has established an initial “guess” parameter set for each mouse that minimizes *ζ*_*i*_, a simplified version of simulated annealing (gradient descent) is performed to refine the optimal parameter prediction. Simulated annealing is a stochastic optimization method with the goal of finding a global optimum [52]. It begins with an initial set of parameters and evolves the parameters with random perturbations until a specified criteria is met. Each new set of randomly perturbed parameters is either accepted or rejected according to the change in *ζ*, which is denoted as 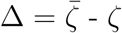 where 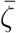 is the SSE of the newly perturbed parameter set and *ζ* is the SSE of last accepted parameter set. If Δ *<* 0 (i.e., the new SSE is lower), then the change is always accepted, and the new parameter set is saved. As numerical experimental revealed that accepting uphill parameter changes decreased algorithm performance (likely due to starting “close to” the optimal from the QMC step), we decided not to accept uphill moves, so if Δ *>* 0, the change not accepted. This algorithm is therefore equivalent to gradient descent. This perturbation process is repeated 5 *×* 10^5^ times for each mouse, and the last accepted parameter set with the lowest *ζ*_*i*_ is taken to be the global optimum. The gradient descent procedure is summarized in Algorithm 2.

#### Algorithm 2: Gradient Descent

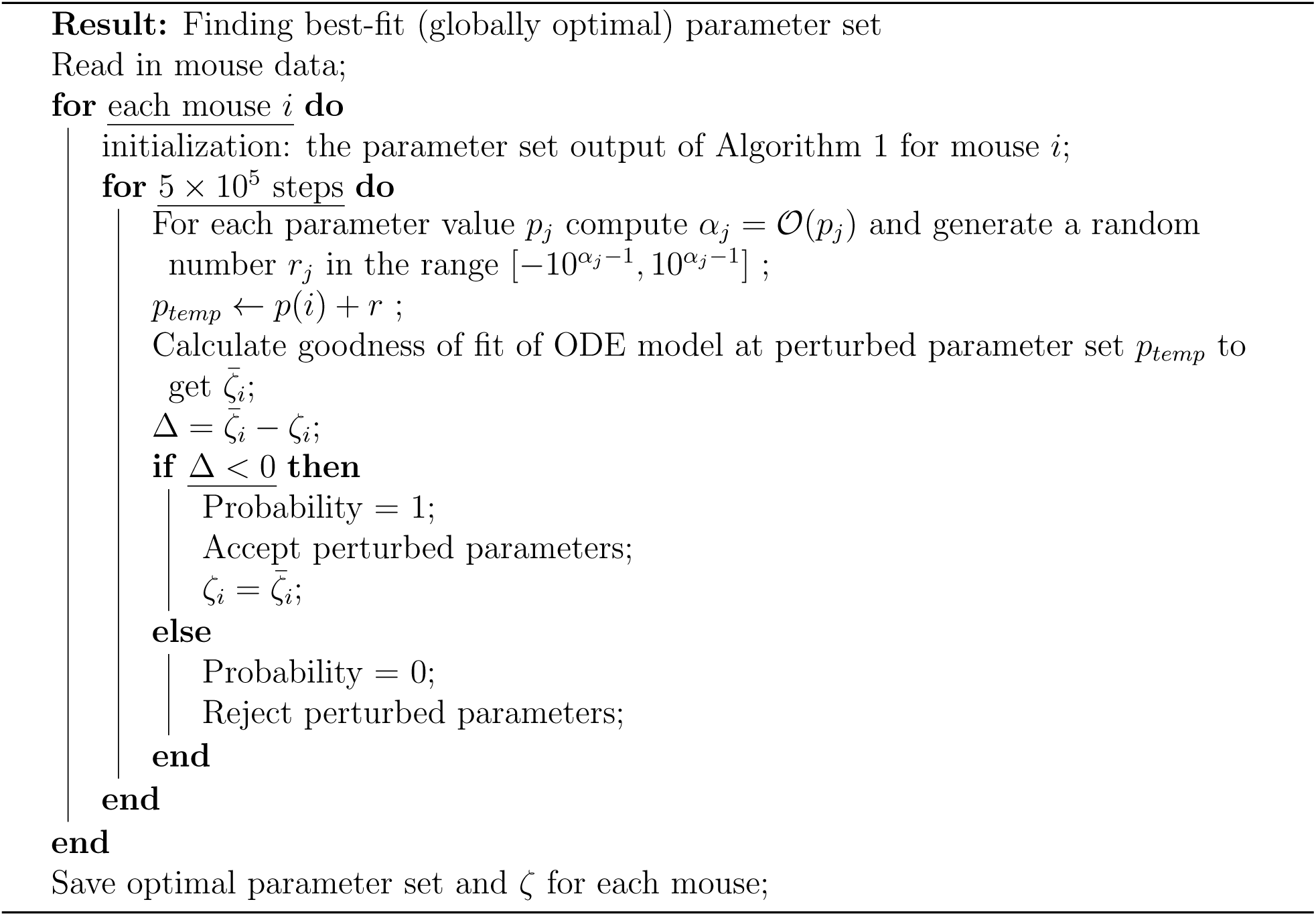

This two-step algorithm is used in all instances, whether fitting the control or treatment data, with the exception of fitting an exponential curve to the control data as that can be done analytically. In all instances where this numerical fitting algorithm was used, the two-step procedure was repeated 15 times, and for each mouse the parameter set with the lowest SSE of the 15 repetitions was chosen as the optimal parameter set.

### 2.5 Identifiability

Identifiability analysis gives us one way to assess the “goodness” of a mathematical model. It is especially relevant in computational models of biological systems given the limited availability and quality (measurement error, noise) of experimental data [54]. Here we will focus on practical identifiability, analyzing if we have sufficient data to have well-determined values for the model parameters. We use profile likelihood (without a confidence interval, since these are fits to the individual, not the average) to evaluate the practical identifiability of a particular model parameter: the initial resistance fraction. This is defined by:

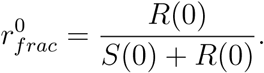

To this end, we define the relevant range for the parameter value, and consider a discrete set of values for the parameter across that range. Since the initial fraction of resistance cells must be in the range [0, 1], we define this range and consider every 0.05 value within this range. At each such 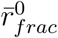 value, we find the best-fit values of the remaining parameters and plot the optimal value of the cost function *ζ*_*i*_ across the parameter’s range to get a profile likelihood curve. The profile likelihood curve for a practically identifiable parameter should appear quadratic, with a clear minimum at the optimal parameter value. If such a quadratic shape is not achieved, the parameter is not practically identifiable [13]. While this analysis could be performed for all model parameters, we choose to focus on the initial resistant fraction, as it is a parameter that is easy to interpret biologically, and therefore will help us in identifying the most likely model describing the experimental data.

## 3 Results and Discussion

To select model that “best” describes the mouse data, we utilized two different model selection methods, the Akaike information criterion (AIC):

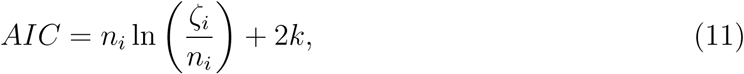

and the Bayesian information criterion (BIC) [38]:

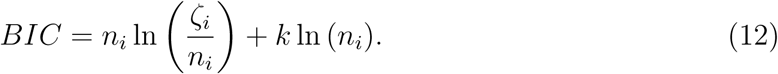

In these equations, *n*_*i*_ is the number of data points for mouse *i, k* is the number of model parameters to be fit, and *ζ*_*i*_ is the SSE from the optimal parameter set for mouse *i*.

Both information criterion consider the trade-offs between goodness of fit (*ζ*_*i*_) and simplicity of the model (i.e. the number of parameters, *k*), allowing us to compare models with different assumptions. However, these measures penalize the number of parameters differently. The AIC model assumes a penalty of the form 2*k*, which is independent of the number of data points. BIC assumes the penalty term *k* ln (*n*_*i*_), meaning the weight of the parameter penalty increases as the number of data points increases. The more data points there are, the greater this penalty term is for the BIC as opposed to the AIC. The model with the lowest AIC (or BIC) score is considered to be the “best”. We will use these two information criterion as we compare our proposed models for the control and treatment data.

### 3.1 Selecting a Model: Control Data

In Section 2.2 we proposed three models to fit the control data: exponential, logistic, and Allee. Mouse 23, 11 and 22 each show different tumor growth behavior of increasing in volume, decreasing in volume, or stabilized volume, respectively. Therefore, we use these three mice in Fig. 2 as representatives to visualize the goodness-of-fit of the various models to the experimental data.

**Figure 2:**
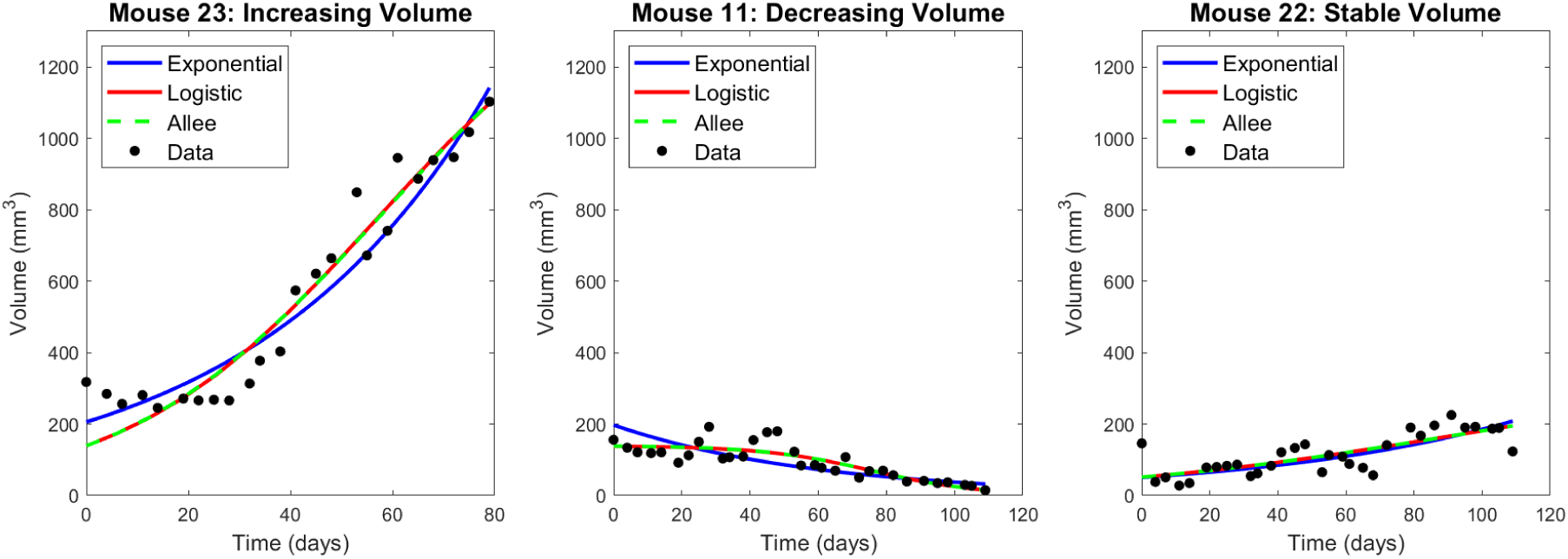
Best fit exponential, logistic, and Allee model for three representative mice. Left (Mouse 23) is representative of the case where the tumor volume increases. Center (Mouse 11) is representative of the case where the tumor volume decreases. Right (Mouse 22) is representative of the case where the tumor volume remains relatively stable.

AIC and BIC values are used for model selection, where the lower the IC value, the better the model is at describing the data. As shown in the top row of Fig. 3, the exponential model appears to be the “best” model, as it has the lowest AIC in 12 of 25 control mice. In 9 of the 12 control mice for which exponential has the lowest AIC, its AIC is at most 5% smaller than the other AIC values for that mouse - we classify this in Fig. 3 as having “low confidence” in the choice for that mouse. The 3 remaining mice for which the exponential model gives the lowest AIC are classified as “medium confidence” (meaning the AIC is 5-10% smaller in best-case). The trends are very similar if we use BIC instead. The exponential has the lowest BIC (and is thus the “best” model) for 14 of the 25 control mice. We have “low confidence” in this prediction for 10 of 13 mice in which exponential has the lowest BIC and “medium confidence” for the remaining 4 mice.

**Figure 3:**
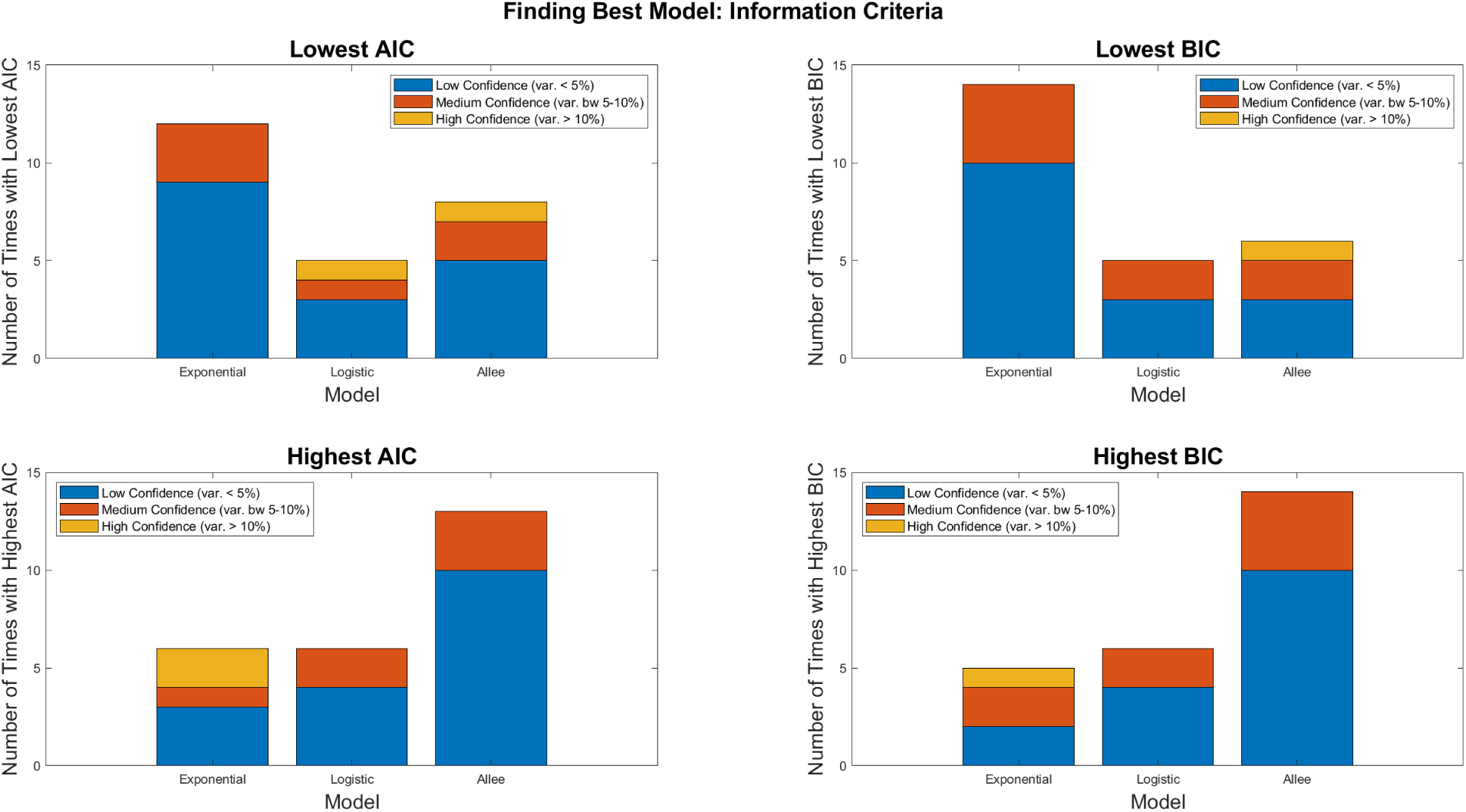
AIC (left column) and BIC (right column) comparisons across control models. Top row shows the number of mice for which each model has the lowest IC value (i.e., is the best model). Bottom row shows the number of mice for which each model has the highest IC value (i.e., is the worst model). We have low confidence in our classification (blue) when the best (or worst, for high IC) IC varies by 5% or less from the other IC values. We have medium confidence (red) when it varies by 5-10%, and high confidence (yellow-orange) when it varies by *>* 10%.

Despite this preference for the exponential model according to both the AIC and BIC, caution is warranted. Selecting a model by minimizing the IC would result in choosing the Allee differential equation for 8 of the mice according to AIC, and 6 of the mice according to BIC. Given the ambiguities in selecting the model for control growth when looking at the lowest IC values, we also looked at the breakdown of which model is the “worst” for describing the data across the control mice; that is, we look for models with the highest IC values. As shown in the bottom row of Fig. 3, the Allee model has the highest IC value in the majority of mice (in 13 of 25 mice according to the AIC, and 14 of 25 mice according to the BIC). Compare this to exponential growth, where the IC is rarely the largest (this occurs in 6 of 25 mice using AIC and 5 of 25 mice using BIC). Considering how often the exponential is the best option of the three (as defined by having the lowest IC) and how rarely it is the worst (as defined by having the highest IC) we will proceed by using an exponential growth term in the treatment data. To assess the robustness, we also consider how our predictions change if we used logistic growth instead.

### 3.2 Insufficiency of volumetric data for treatment model selection

In Section 2.3, we proposed a family of six models to describe the resistance of cetuximab in our xenograft data (see Fig. 1): Model 1.1 with no resistance, Model 1.2 with pre-existing resistance only, Model 2.1 with randomly-acquired resistance only, Model 2.2 with randomly-acquired and pre-existing resistance, Model 3.1 with drug-induced resistance only, and Model 3.2 with drug-induced and pre-existing resistance. Assuming exponential growth as justified in Section 3.1, the fits of the six treatment models to data for three representative mice that exhibit different tumor growth dynamics are shown in Fig. 4. The best-fit value of all parameters, across all mice and models, is shown in Fig. S2. An interesting observation is that whenever the model allows for pre-existing resistance, resistant cells are present in substantial numbers prior to treatment. In particular, Model 1.2 has a median resistance fraction of 53.01%, with a mean of 57.18 ± 27.28%. Model 2.2 has a median resistance fraction of 25.55%, with a mean of 35.96 ± 28.80%. Model 3.2 has a median initial resistance fraction of 10.48%, with a mean of 31.75 ± 38.06%. While these values do seem large when considering that resistance often results in a fitness disadvantage in the absence of drug [47, 6], the existence of a significant pool of HNSCC resistant to cetuximab prior to treatment is consistent with single-cell data from cetuximab sensitive HNSCC cell lines [29].

**Figure 4:**
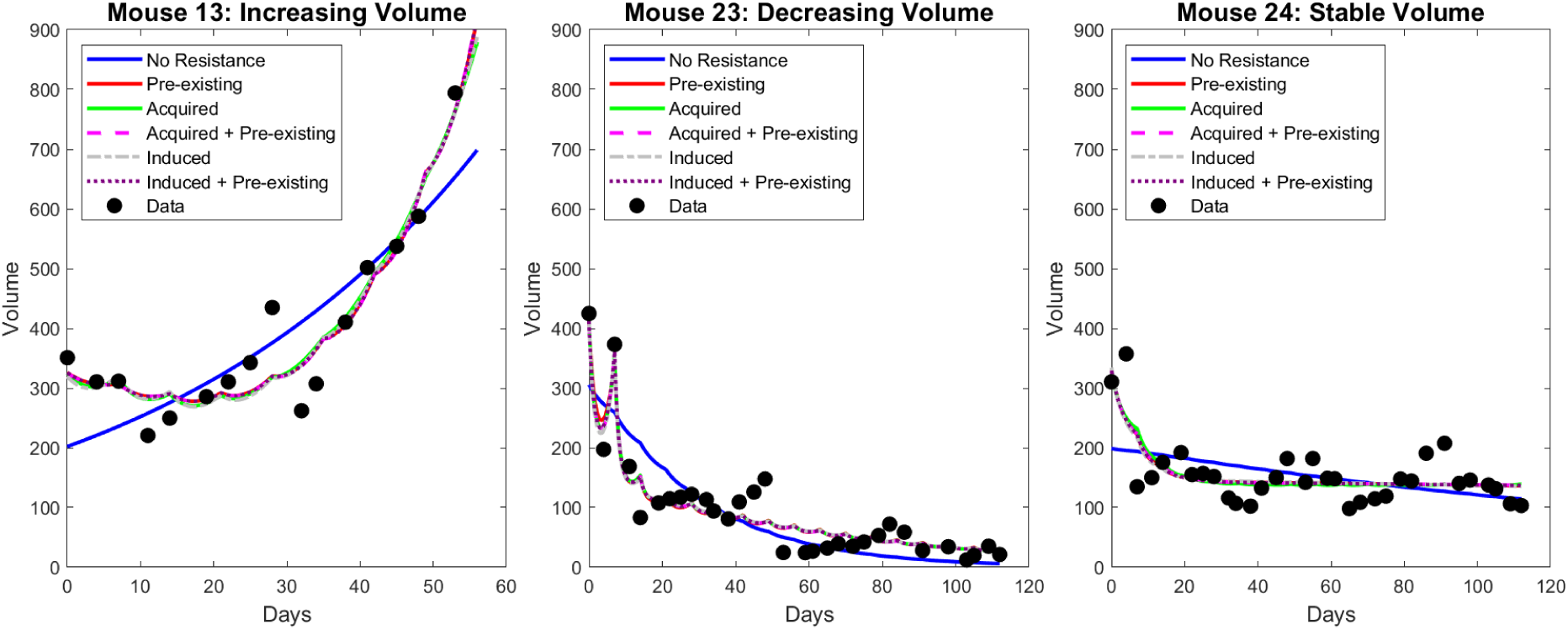
Best fit of six proposed resistance models to treatment data for three representative mice. Left (Mouse 13) is representative of the case where the tumor volume increases in spite of treatment. Center (Mouse 23) is representative of the case where the tumor volume decreases during treatment. Right (Mouse 24) is representative of the case where the tumor volume remains relatively stable during treatment.

A visual inspection alone suggests that Model 1.1 (no resistance) cannot adequately explain treatment response to cetuximab. In order to quantitatively approach model selection so as to determine which “member(s)” of our family of resistance models most likely captures the mechanisms in the data, we computed the AIC and BIC for each mouse and model (Fig. 5). This analysis confirms that some form of resistance must be driving treatment response to cetuximab, as Model 1.1 very rarely has the lowest IC value (happens 2 of 29 times for AIC, and 4 of 29 times for BIC), and very frequently has the highest IC value (happens 22 of 29 times for AIC, and 19 of 29 times for BIC). By a similar argument, the IC values indicate that the resistance in the data likely cannot be attributed to a combination of randomly acquired and pre-existing resistance (Model 2.2). As indicated in Fig. 5, Model 2.2 never has the lowest IC value, and occasionally has the highest IC value (happens 4 of 29 times for AIC, and 9 of 29 times for BIC). Therefore, this information theoretic analysis was able to rule out several mechanistic explanations of cetuximab resistance, but it is not sufficient to select the model whose mechanisms most likely explain this resistance. Notably, the case of no resistance, and the case of resistance being pre-existing and randomly-acquired, are also ruled out if growth is assumed to be logistic instead of exponential (see Fig. S3).

**Figure 5:**
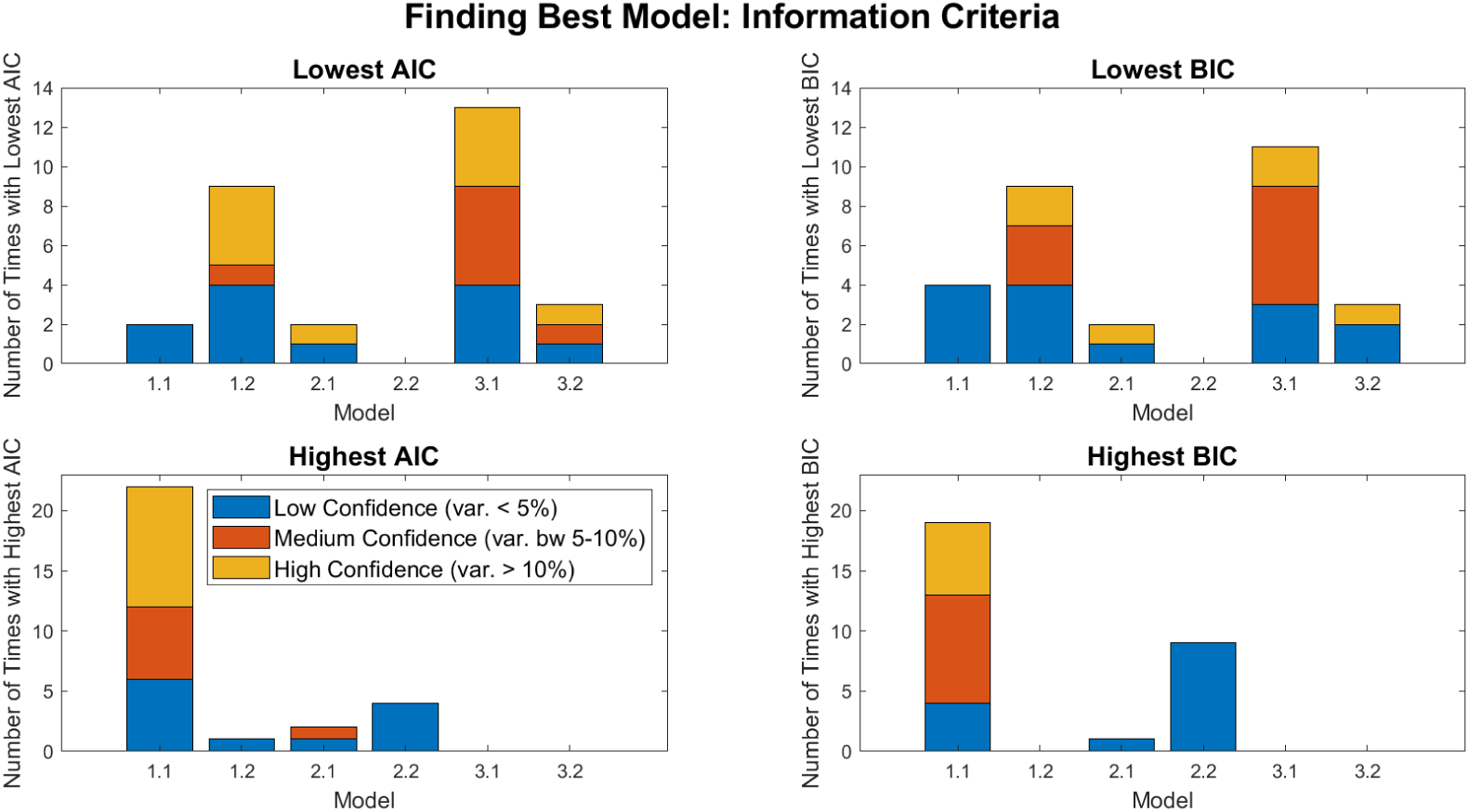
AIC (left column) and BIC (right column) comparisons across treatment models when exponential growth is used. Top row shows the number of mice for which each model has the lowest IC value (i.e., is the best model). Bottom row shows the number of mice for which each model has the highest IC value (i.e., is the worst model).

### 3.3 Initial pre-existing resistance fraction facilitates treatment model selection

Our analyses thus far assume that the only available data is the time-course describing tumor volume in individual xenografts. However, advances in single-cell technology now allows the initial fraction of resistant cells in a tumor to be quantified [29]. While such analyses were not undertaken for the volumetric data presented herein, our modeling framework could be readily used to ask: does the inclusion of the initial resistance fraction improve our model selection capabilities? Trivially, knowledge that there are or are not pre-existing resistant cells automatically eliminates half the models considered in our family of models. Therefore, for the sake of this analysis, we will assume that resistance cells exist prior to treatment, as demonstrated in [29]. Under this assumption we now explore how having this additional data point does, or does not, facilitate treatment model selection.

We approach this question by determining the practical identifiability of the initial resistance fraction in our models using profile likelihood. We consider Model 1.2 (pre-existing resistance only) and Model 3.2 (drug-induced acquired plus pre-existing resistance), though not Model 2.2 as our information theoretic analysis already concluded that the combination of randomly acquired and pre-existing resistance is a highly unlikely to explain cetuximab resistance in our xenograft data. In 20 of 29 mice fit using Model 1.2, the profile likelihood curves for the initial resistance fraction reveal this parameter to be practically identifiable. The profile likelihood curve for Mouse 1 (left panel of Fig. 6) is representative of these 20 mice. For Mouse 1, we observe a clear optimal value for the initial resistance at 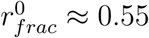. Any significant deviation from this resistance fraction drastically increases the cost function (that is, decreases the goodness-of-fit). In this case, having the true value of the initial resistance fraction could greatly inform the process of model selection. If the true initial resistance fraction was 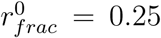, the cost function increases three-fold. This in turn increases the IC values in eqns. (11) and (12), and significantly reduces the likelihood of Model 1.2 having the lowest IC values. In other words, this would provide strong evidence that pre-existing resistance alone does not explain cetuximab resistance in the data.

**Figure 6:**
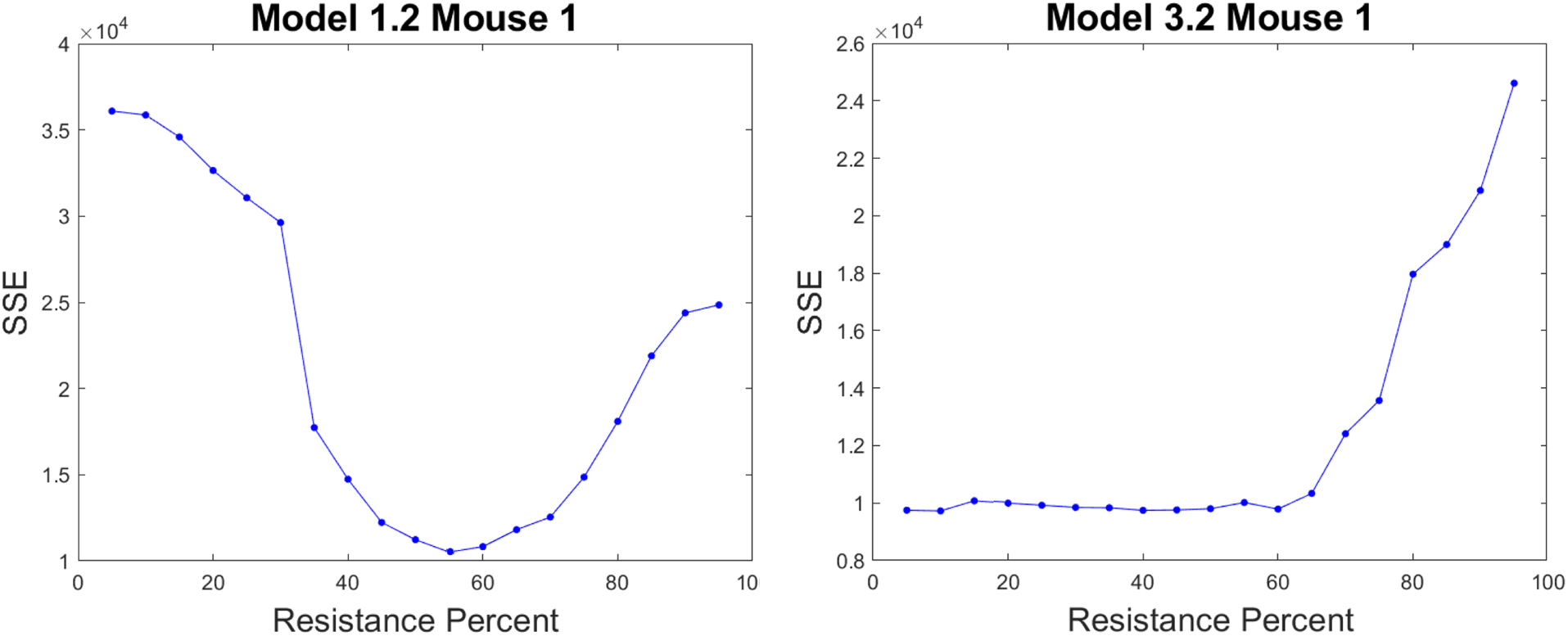
Profile likelihood curves of the initial resistance fraction for a representative mouse, Mouse 1. Left curve (Model 1.2) is practically identifiable and places the optimal parameter at approximately 55%. Right curve (Model 3.2) is not practically identifiable.

Compare this to what happens in the same mouse fit using Model 3.2 (pre-existing plus drug-induced resistance). The profile likelihood curve for the same parameter is not practically identifiable, as demonstrated in Fig. 6 (right panel) by a shallow profile with a one-sided minimum. This is not unique to Mouse 1 - the profile likelihood curves for the initial resistance fraction in Model 3.2 reveal this parameter to be practically non-identifiable in 22 of 29 mice. Returning to our prior thought experiment, for Mouse 1 in particular, if we had measured the initial resistance fraction to be 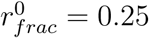, we would have strong evidence that Model 3.2 should be selected over Model 1.2. While the lack of practical identifiability poses mathematical challenges, it does give Model 3.2 a lot more “flexibility” to conform to additional experimental data without sacrificing goodness-of-fit.

Finally, it is important to note that the addition of the initial resistance fraction is not always sufficient to select a model. Continuing to use Mouse 1 as an example, if we had measured the true initial resistance fraction to be 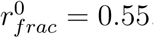, both Models 1.2 and 3.2 remain viable choices. Therefore we conclude that while measuring the initial resistance fraction is an essential step to understanding the mechanisms underlying cetuximab resistance, it is not guaranteed to determine the mechanism of resistance using our family of mathematical models.

### 3.4 Dose escalation study further facilitates treatment model section

Thus far we have established that time-course volumetric data combined with a measurement of the initial resistance fraction may or may not be sufficient to deduce the underlying mechanism of resistance using our family of models. Here, we propose a final experiment that, combined with the other data, would be sufficient to select a treatment model. In particular, we propose a dose escalation study where we use the optimal parameter set for each mouse to simulate tumor response to a range of drug doses. We measure the fold reduction in the tumor volume per mouse by comparing the initial tumor volume in each mouse to its volume two weeks later, with one dose of cetuximab given per week as in the experimental protocol. The median fold reduction across all 29 mice, at each dose, is then computed (see Fig. 7). As an example, a median fold reduction of 4 means the median tumor volume is four times smaller post-treatment than it was pre-treatment. Thus a higher median fold reduction represents a more effective treatment.

**Figure 7:**
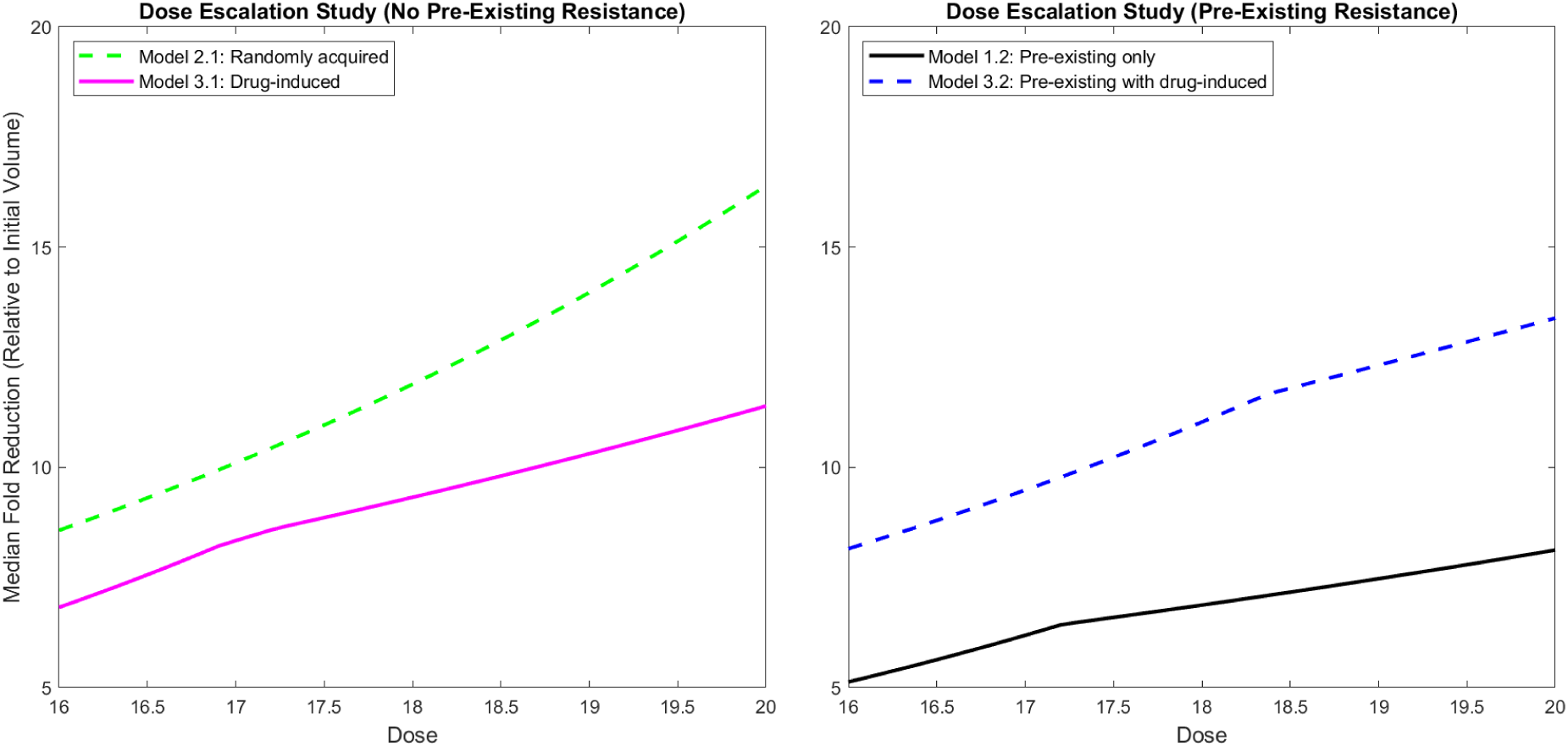
Dose escalation study of median reduction in tumor volume (relative to initial volume) after 2 weeks. Dose varies from 16 to 20 mg/kg. Growth is assumed to be exponential. Left: plausible models involving no pre-existing resistance. Right: plausible models including pre-existing resistance.

As shown in the left panel of Fig. 7, in the case of no pre-existing resistance, the two plausible models show drastically different responses to a dose escalation. Starting at a dose of 16 mg/kg, simulations show a much more significant fold reduction in median tumor volume when resistance is randomly acquired (8.571 median fold reduction) than when resistance is drug-induced (6.822 median fold reduction). The lower response in the drug-induced case can be explained by the fact that the transition from the sensitive to the resistant phenotype is directly promoted by the drug itself. Thus higher drug doses drive more resistance formation in the drug-induced case, though not in the randomly acquired case.

Looking across doses, we also observe a noticeably different change in the median volumetric fold-reduction as the dose is escalated from 16 mg/kg to 20 mg/kg. The average rate of change in the randomly acquired case is 1.955, whereas the average rate of change in the drug-induced case is only 1.143. This strongly suggests that one way to distinguish between modes of acquired resistance, at least in the absence of pre-existing resistance, is to experimentally perform this dose escalation study.

The right panel of Fig. 7 shows the two plausible models in the case of pre-existing resistance. Focusing on the dose of 16 mg/kg, simulations show a more significant fold reduction in median tumor volume when resistance can be induced by the drug. This occurs because the initial resistance fraction required to fit the data when the model does not include drug-induced resistance is necessarily larger than the initial resistance fraction when there is a secondary mechanism for creating resistant cells (in the case of Model 3.2, the drug itself promotes the transition to resistance). Therefore, Model 1.2 mice always have a larger initial resistance fraction, and thus experience a smaller response to the drug over the relatively short time period of two weeks. However, as observed in the case of no pre-existing resistance, we still observe a more significant change in the median fold reduction in the case of drug-induced resistance as the dose is escalated from 16 mg/kg to 20 mg/kg. In particular, the average rate of change in the drug-induced case is 1.309, whereas the average rate of change in the case where all resistance is pre-existing is 0.749. Whether resistance is pre-existing or not, we see an approximately 1.7-fold difference in the median across models, demonstrating that an experimental dose escalation study would provide meaningful data in trying to elucidate the mechanisms driving cetuximab resistance.

## 4 Conclusion

In this work, we introduced a family of six ordinary differential equation models, with each model assuming a different underlying biological mechanism(s) driving cetuximab resistance in patient-derived xenografts of head and neck squamous cell carcinoma. Model selection techniques alone allowed us to conclude that some form of resistance must be driving the treatment response dynamics, and that this resistance was highly unlikely to be explained by randomly acquired resistance coupled with pre-existing resistance.

With four remaining family members remaining to plausibly describe cetuximab resistance, we next asked: what additional data would be needed so that we can identify model from the family that is most parsimonious with the data? Through the use of profile likelihood curves, we uncover that quantifying the initial fraction of resistant cells in a tumor population, which can now be readily done due to advances in single-cell technology [29], improves the likelihood of identifying the model within the family that is most parsimonious with the data. Finally, we find that if this experiment is coupled with a dose escalation study, the combination of the experimental data and mathematical modeling allows the mechanism and timing of cetuximab resistance to be determined.

The conclusions drawn in this work are dependent on the family of models constructed. While we have demonstrated some robustness in the results to the underlying growth term in the absence of treatment, other functional forms for both tumor growth and drug effects could certainly be considered. As future work, one option would be to expand the family of models and repeating the analyses herein to determine if a combination of time-course volumetric data at a single dose, measurements of the initial resistance fraction, and dose escalation data are sufficient experimental data to pinpoint the mechanism and timing of resistance. Alternatively, model learning techniques [4, 11, 8, 36, 55, 24, 42, 44], possibly informed by biological knowledge [32], provide tools for considering a much larger class of mathematical models of cetuximab resistance in HNSCC.

Understanding the mechanisms driving cetuximab resistance is essential, as optimal therapeutic design is likely dependent on the underlying mode(s) of resistance. For instance, work in [23] computationally demonstrated that tumor response to the same drug dose and delivery schedule is qualitatively impacted by the ability (or lack therefore) of a drug to induce resistance. Therefore it is essential that any mathematical model accurately capture the mechanism driving resistance if that model is to be used to optimize drug dosing and the delivery schedule. This work demonstrates how mathematical modeling can inform future experimental design, allowing for an improved understanding of dosing novel cancer therapeutics.

## Data Availability

All experimental data and fitting codes are publicly available at https://github.com/jgevertz/HNSCC-Cetuximab-Resistance.

## Acknowledgments

The authors thank Luciane T. Kagohara, Daria A. Gaykalova, and Genevieve Stein-O’Brien for feedback on this study. JLG, SDC, and CR acknowledge use of the ELSA high-performance computing cluster at The College of New Jersey for conducting the research reported in this paper. This cluster is funded in part by the National Science Foundation under grant numbers OAC-1826915 and OAC-1828163. Funding was obtained from the National Institutes of Health National Cancer Institute (R01CA177669) and the Johns Hopkins University Catalyst Award to EJF.

## Author Contributions

All authors substantially contributed to the either the collection or analysis the data in this work, and the drafting of the manuscript. Each author also provides approval for publication of the content. RR, FS, and CHC performed the patient-derived xenograft studies. SDC, CR, and JLG performed the mathematical modeling. EJF and JLG are responsible for the design of the work.

## Competing Interests

The authors declare the following competing interests: EJF is on the Scientific Advisory Board of ResistanceBio/Viosera Therapeutics and is a paid consultant to Merck and Mestag Therapeutics. CHC has honoraria from Bristol-Myers Squibb, CUE, Sanofi, Mirati, Merck, Brooklyn ImmunoTherapuetics, and Exelixis for ad hoc Scientific Advisory Board participation. RR was affiliated with Moffitt at the time of data generation, but employed by BMS at time of submission.

## 5 Supplemental Information

**Table S1:** Volume measurements (mm^3^) for the 25 control mice and 29 treatment mice. Note that Mouse *m* in the control case is not related to Mouse *m* in the treatment case.

| Mouse ID | Initial Control Volume (mm<sup>3</sup>) | Initial Treatment Volume (mm<sup>3</sup>) |
| --- | --- | --- |
| 1 | 150 | 538.28 |
| 2 | 473.8 | 298.58 |
| 3 | 549.5 | 159.83 |
| 4 | 596.8 | 288.71 |
| 5 | 346.4 | 871.67 |
| 6 | 522.6 | 501.84 |
| 7 | 571.8 | 105.47 |
| 8 | 1008.4 | 279.4 |
| 9 | 442 | 410.60 |
| 10 | 628.6 | 279.20 |
| 11 | 155.9 | 547.97 |
| 12 | 339.8 | 205.27 |
| 13 | 624.4 | 350.98 |
| 14 | 1073.2 | 616.01 |
| 15 | 559.4 | 1188.23 |
| 16 | 281 | 524.62 |
| 17 | 727.8 | 935.25 |
| 18 | 410.6 | 391.13 |
| 19 | 550.1 | 551.35 |
| 20 | 422.2 | 1274.23 |
| 21 | 909.6 | 452.07 |
| 22 | 145.8 | 348.55 |
| 23 | 317.7 | 424.87 |
| 24 | 228.3 | 310.44 |
| 25 | 724.9 | 744.675 |
| 26 | — | 397.37 |
| 27 | — | 1111.57 |
| 28 | — | 1097.87 |
| 29 | — | 842.41 |

**Figure S1:**
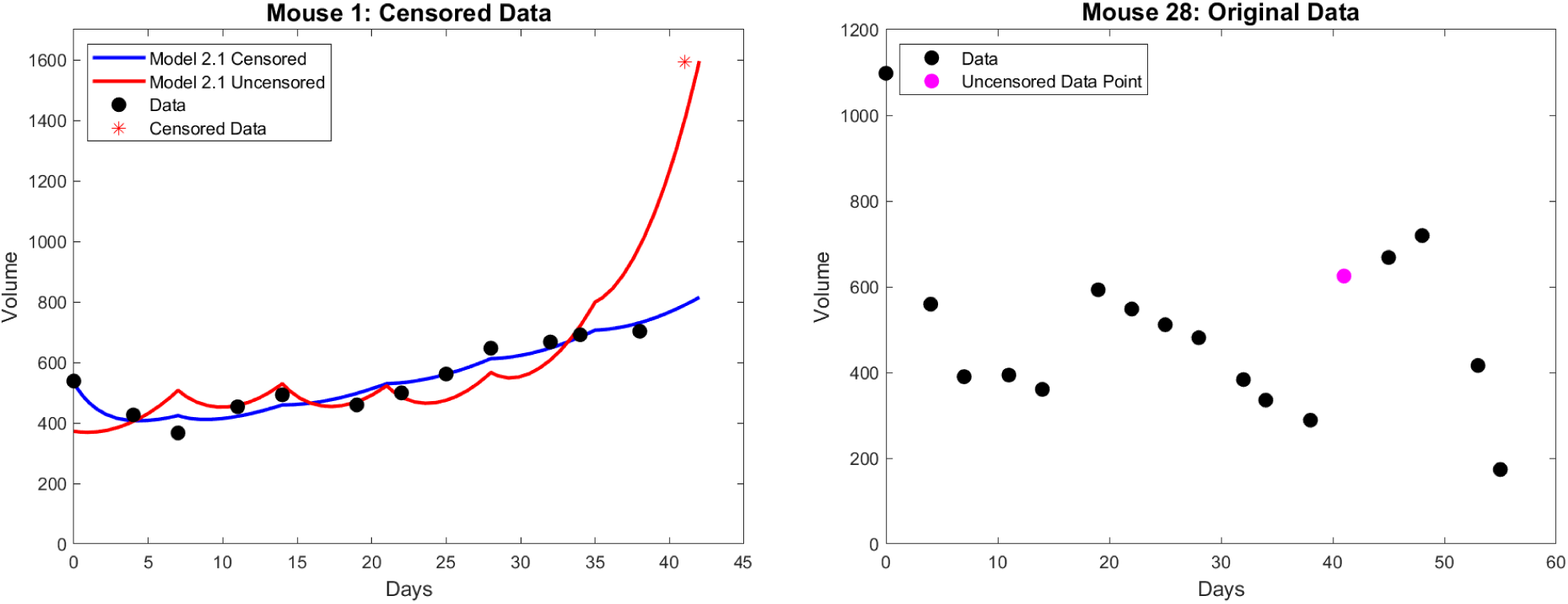
Two representative treatment mice to depict censoring protocol. Uncensored data for Mouse 1 (left panel) is shown in black dots with the censored point as a red asterisk. The best-fit curve from Model 1.2 on both censored and uncensored data is shown. Mouse 28 (right panel) had no censored data points. The magenta data point is candidate for censoring because of the rapid growth over a short period of time, but it is not censored because the subsequent points follow the trend.

**Figure S2:**
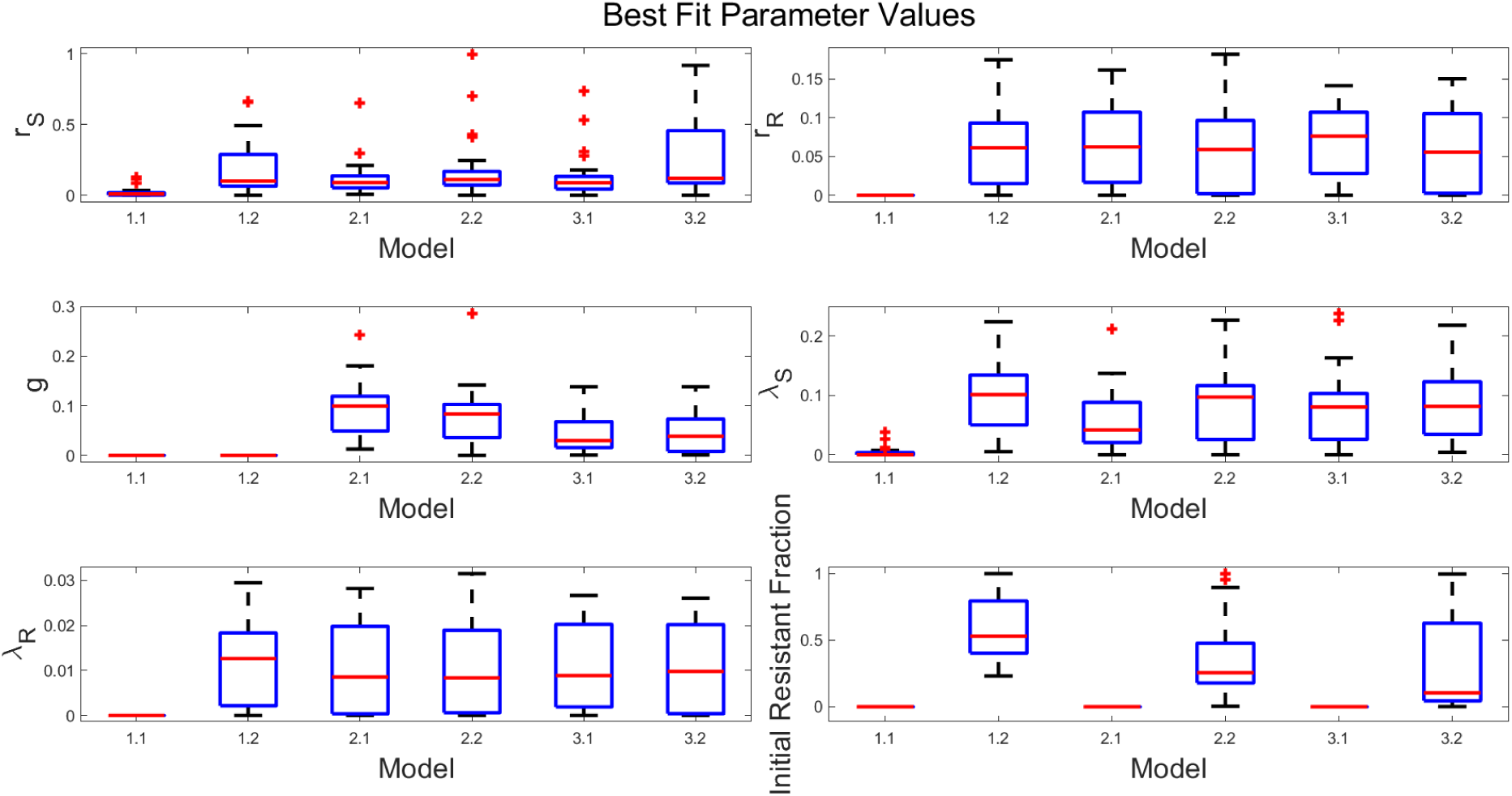
Box plot showing best-fit parameter values across six models for sensitive cell growth rate *r*_*S*_ (top left), resistant cell growth rate *r*_*R*_ (top right), rate of random transition to resistance *g* (center left), drug-induced death rate of sensitive cells *λ*_*S*_ (center right), drug-induced death rate of resistant cells *λ*_*R*_ (bottom left), and initial resistant fraction 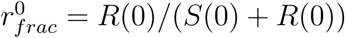 (bottom right).

**Figure S3:**
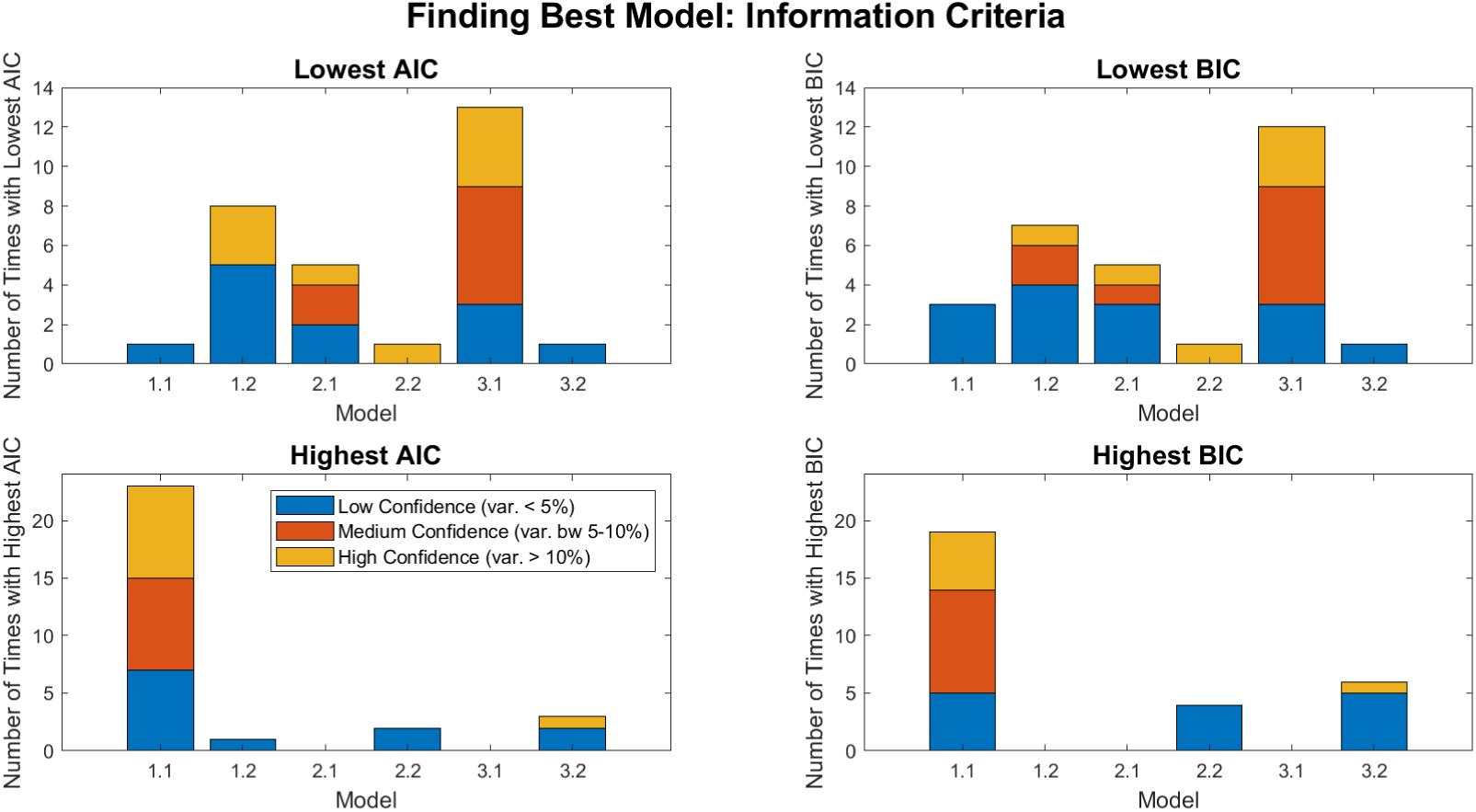
AIC (left column) and BIC (right column) comparisons across treatment models when logistic growth is used. Top row shows the number of mice for which each model has the lowest IC value (i.e., is the best model). Bottom row shows the number of mice for which each model has the highest IC value (i.e., is the worst model).

## Notes

https://github.com/jgevertz/HNSCC-Cetuximab-Resistance

## References

[1] Erbitux. ImClone Systems Incorporated and Bristol-Meyers Squibb Company. URL: https://www.accessdata.fda.gov/drugsatfda_docs/label/2004/125084lbl.pdf.

[2] D.E. Axelrod, S. Vedula, and J. Obaniyi. Effective chemotherapy of heterogeneous and drug-resistant early colon cancers by intermittent dose schedules: a computer simulation study. Cancer Chemother. Pharmocol., 79:889–898, 2017.

[3] C.C. Bell and O. Gilan. Principles and mechanisms of non-genetic resistance in cancer. Br. J. Cancer, 122:465–472, 2020.

[4] Josh Bongard and Hod Lipson. Automated reverse engineering of nonlin-ear dynamical systems. Proceedings of the National Academy of Sciences, 104(24):9943–9948, 2007. URL: https://www.pnas.org/content/104/24/9943 http://arxiv.org/abs/https://www.pnas.org/content/104/24/9943.full.pdf arXiv:https://www.pnas.org/content/104/24/9943.full.pdf, https://doi.org/10.1073/pnas.0609476104 doi:10.1073/pnas.0609476104.

[5] J.A. Bonner, P.M. Harari, J. Giralt, N. Azarnia, D.M. Shin, R.B. Cohen, Jones C.U., R. Sur, D. Raben, J. Jassem, R. Ove, M.S. Kies, J. Baselga, H. Youssoufian, N. Amellal, E.K. Rowinsky, and K.K. Ang. Radiotherapy plus cetuximab for squamous-cell carcinoma of the head and neck. N Engl J Med., 354:567–578, 2006.

[6] K.R. Brimacombe, M.D. Hall, and et. al. Auld, D.S. A dual-fluorescence high-throughput cell line system for probing multidrug resistance. Assay Drug Dev. Technol., 7:233–249, 2009.

[7] T. Brocato, P. Dogra, E. J. Koay, A. Day, Y-L. Chuang, Z. Wang, and V. Cristini. Understanding drug resistance in breast cancer with mathematical oncology. Curr. Breast Cancer Rep., 6:110–120, 2014.

[8] Steven L. Brunton, Joshua L. Proctor, and J. Nathan Kutz. Discovering governing equations from data by sparse identification of nonlinear dynamical systems. Proceedings of the National Academy of Sciences, 113(15):3932–3937, 2016. URL: https://www.pnas.org/content/113/15/3932, http://arxiv.org/abs/https://www.pnas.org/content/113/15/3932.full.pdf arXiv:https://www.pnas.org/content/113/15/3932.full.pdf, https://doi.org/10.1073/pnas.1517384113 doi:10.1073/pnas.1517384113.

[9] M.P. Chapman, T.T. Risom, A. Aswani, R. Dobbe, R.C. Sears, and C.J. Tomlin. A model of phenotypic state dynamics initiates a promising approach to control heterogeneous malignant cell populations. In 2016 IEEE 55th Conference on Decision and Control (CDC), pages 2481–2487. IEEE, December 2016.

[10] R.H. Chisholm, T. Lorenzi, A. Lorz, A.K. Larsen, L. Neves de Almedia, A. Escargueil, and J. Clairambault. Emergence of drug tolerance in cancer cell populations: an evolutionary outcome of selection, nongenetic instability, and stress-induced adaptation. Cancer Research, 75:930–939, 2015.

[11] B. Daniels and I. Nemenman. Automated adaptive inference of phenomenological dynamical models. Nat. Commun., 6:8133, 2015.

[12] H. Du, Y. Zhao, H. Li, D.W. Wang, and C. Chen. Roles of micrornas in glucose and lipid metabolism in the heart. Front Cardiovasc Med., 8:716213, 2021.

[13] Marisa C. Eisenberg and Harsh V. Jain. A confidence building exercise in data and identifiability: Modeling cancer chemotherapy as a case study. J Theor Biol., 431:63– 78, 2017.

[14] M.S. Feizabadi. Modeling multi-mutation and drug resistance: analysis of some cases. Theoretical Biology and Medical Modelling, 14:6, 2017.

[15] J. Foo, K. Leder, and S.M. Mumenthaler. Cancer as a moving target: understanding the composition and rebound growth kinetics of recurrent tumors. Evol. Appl., 6:54–69, 2013.

[16] J. Foo and F. Michor. Evolution of resistance to targeted anti-cancer therapies during continuous and pulsed administration strategies. PLoS Comput. Biol., 5:e1000557, 2009.

[17] J. Foo and F. Michor. Evolution of acquired resistance to anti-cancer therapy. J. Theor. Biol., 355:10–20, 2014.

[18] A. A. Forastiere, M. Maor, R. S. Weber, T. Pajak, A. Trotti, J. Ridge, J. Ensley, C. Chao, and J. Cooper. Long-term results of intergroup rtog 91-11: A phase iii trial to preserve the larynx-induction cisplatin/5-fu and radiation therapy versus concurrent cisplatin and radiation therapy versus radiation therapy. Journal of Clinical Oncology, 24, 2006.

[19] Jake C. Forster, Michael J. J. Douglass, Wendy M. Harriss-Phillips, and Eva Bezak. Simulation of head and neck cancer oxygenation and doubling time in a 4d cellular model with angiogenesis. Sci Rep., 7:11037, 2017.

[20] J.L. Gevertz, Z. Aminzare, K.-A. Norton, J. Perez-Velazquez, A. Volkening, and K.A. Rejniak. Emergence of anti-cancer drug resistance: exploring the importance of the microenvironmental niche via a spatial model. In T. Jackson and A. Radunskaya, editors, Applications of Dynamical Systems in Biology and Medicine, volume 158 of The IMA Volumes in Mathematics and its Applications, pages 1–34. Springer-Verlag, 2015.

[21] A. Goldman, B. Majumder, A. Dhawan, S. Ravi, D. Goldman, M. Kohandel, P.K. Majumder, and S. Sengupta. Temporally sequenced anticancer drugs overcome adaptive resistance by targeting a vulnerable chemotherapy-induced phenotypic transition. Nature Communications, 6:6139, 2015.

[22] J. Greene, O. Lavi, M.M. Gottesman, and D. Levy. The impact of cell density and mutations in a model of multidrug resistance in solid tumors. Bull. Math. Biol., 76:627– 653, 2014.

[23] James M. Greene, Jana L. Gevertz, and Eduardo D. Sontag. Mathematical approach to differentiate spontaneous and induced evolution to drug resistance during cancer treatment. JCO Clinical Cancer Informatics, 3:1–20, 2019.

[24] Roger Guimerà, Ignasi Reichardt, Antoni Aguilar-Mogas, Francesco A. Massucci, Manuel Miranda, Jordi Pallarès, and Marta Sales-Pardo. A bayesian machine scientist to aid in the solution of challenging scientific problems. Science Advances, 6(5):eaav6971, 2020.

[25] Vanessa Hearnden, Hilary J. Powers, Abeir Elmogassabi, Rosanna Lowe, and Craig Murdoch. Methyl-donor depletion of head and neck cancer cells in vitro estabilishes a less aggressive tumour cell phenotype. Eur J Nutr., 57:1321–1332, 2018.

[26] T.L. Jackson and H.M. Byrne. A mathematical model to study the effects of drug resistance and vasculature on the response of solid tumors to chemotherapy. Math. Biosci., 164:17–38, 2000.

[27] K.E. Johnson, G. Howard, W. Mo, M.K. Strasser, E.A.B.F. Lima, S. Huang, and A. Brock. Cancer cell population growth kinetics at low densities deviate from the exponential growth model and suggest an Allee effect. PLoS Biol., 17:e3000399, 2019.

[28] M.K. Jolly, P. Kulkarni, K Weninger, J. Orban, and H. Levine. Phenotypic plasticity, bet-hedging, and androgen independence in prostate cancer: Role of non-genetic heterogeneity. Front. Oncol., 8:50: doi: 10.3389/fonc.2018.00050, 2018.

[29] F. Kagohara, L.T. amd Zamuner, E.F. Davis-Marcisak, G. Sharma, M. Considine, J. Allen, S. Yegnasubramanian, D.A. Gaykalova, and E.J. Fertig. Integrated single-cell and bulk gene expression and atac-seq reveals heterogeneity and early changes in pathways associated with resistance to cetuximab in hnscc-sensitive cell lines. Br J Cancer., 123:1582–1583, 2020.

[30] N.L. Komarova and D. Wodarz. Drug resistance in cancer: Principles of emergence and prevention. Proc. Natl. Acad. Sci., 102:9714–9719, 2005.

[31] Sergei Kucherenko, Daniel Albrecht, and Andrea Saltelli. Exploring multi-dimensional spaces: a comparison of latin hypercube and quasi monte carlo sampling techniques. arXiv Cornell University, 2015.

[32] J.H. Lagergren, J.T. Nardini, R.E. Baker, M.J. Simpson, and K.B. Flores. Biologically-informed neural networks guide mechanistic modeling from sparse experimental data. PLoS Comput Biol., 16:e1008462, 2020.

[33] O. Lavi, M.M. Gottesman, and D. Levy. The dynamics of drug resistance: A mathematical perspective. Drug Resist. Update, 15:90–97, 2012.

[34] Quynh-Thu Le and David Raben. Integrating biologically targeted therapy in head and neck squamous cell carcinomas. Seminars in radiation oncology, 19(1):53—62, January 2009. https://doi.org/10.1016/j.semradonc.2008.09.010 doi:10.1016/j.semradonc.2008.09.010.

[35] L.L. Liu, F. Li, W. Pao, and F. Michor. Dose-dependent mutation rates determine optimum erlotinib dosing strategies for EGRF mutant non-small lung cancer patients. PLoS ONE, 10:e0141665, 2015.

[36] N.M. Mangan, J.N. Kutz, S.L. Brunton, and J.L. Proctor. Model selection for dynamical systems via sparse regression and information criteria. Proc. R. Soc. A, 473:20170009, 2017.

[37] Ranee Mehra, Roger B. Cohen, and Barbara A. Burtness. The role of cetuximab for the treatment of squamous cell carcinoma of the head and neck. PubMed Central, 10:742–750, 2009.

[38] H. Miao, C. Dykes, L Demeter, and W. Hulin. Differential equation modeling of hiv viral fitness experiments: model identification, model selection, and multimodel inference. Biometrics, 65:292–300, 2009.

[39] M. Mrhalova, J. Plzak, J. Betka, and R. Kodet. Epidermal growth factor receptor–its expression and copy numbers of egfr gene in patients with head and neck squamous cell carcinomas. Neoplasma, 52:338–343, 2005.

[40] S.M. Mumenthaler, J. Foo, K. Leder, N.C. Choi, D.B. Agus, W. Pao, P. Mallick, and F. Michor. Evolutionary modeling of combination treatment strategies to overcome resistance to tyrosine kinase inhibitors in non-small cell lung cancer. Mol. Pharm., 8:2069–2079, 2011.

[41] Hope Murphy, Hana Jaafari, and Hana M Dobrovolny. Differences in prediction of ode models of tumor growth: a cautionary example. BMC Cancer, 16:1–10, 2016.

[42] J.T. Nardini, J.H. Lagergren, A Hawkins-Daarud, L. Curtin, B. Morris, E.M. Rutter, K.R. Swanson, and K.B. Flores. Learning Equations from Biological Data with Limited Time Samples. Bull. Math. Biol., 82:119, 2020.

[43] Angela Oliveira Pisco, Amy Brock, Joseph Zhou, Andreas Moor, Mitra Mojtahedi, Dean Jackson, and Sui Huang. Non-darwinian dynamics in therapy-induced cancer drug resistance. Nature communications, 4, 2013.

[44] H. Ribera, S. Shirman, A.V. Nguyen, and N.M. Mangan. Model selection of chaotic systems from data with hidden variables using sparse data assimilation. arXiv Cornell University, 2021.

[45] R.T. Santuray, D.E. Johnson, and J.R. Grandis. New therapies in head and neck cancer. Trends Cancer, 4:385–396, 2018.

[46] E.A. Sarapata and L.G. de Pillis. A comparison and catalog of intrinsic tumor growth models. Bull. Math. Biol., 76:2010–2024, 2019.

[47] S.E. Shackney, G.W. McCormack, and G.J. Jr Cuchural. Growth rate patterns of solid tumors and their relation to responsiveness to therapy: An analytical review. Ann. Intern. Med., 89:107–121, 1978.

[48] S.V. Sharma, D.Y. Lee, B. Li, M.P. Quinlan, F. Takahashi, S. Maheswaran, U. McDermott, N Azizian, L. Zou, M.A. Fischbach, K.K. Wong, K. Brandstetter, B. Wittner, S. Ramaswamy, M. Classon, and J. Settleman. A chromatin-mediated reversible drug-tolerant state in cancer cell subpopulations. Front. Oncol., 141:69–80, 2010.

[49] A.S. Silva and R.A. Gatenby. A theoretical quantitative model for evolution of cancer chemotherapy resistance. Biol. Direct., 5:25, 2010.

[50] Genevieve Stein-O’Brien, Luciane T. Kagohara, Sijia Li, Manjusha Thakar, Ruchira Ranaweera, Hiroyuki Ozawa, Haixia Cheng, Michael Considine, Sandra Schmitz, Alexander V. Favorov, Ludmila V. Danilova, Joseph A. Califano, Evgeny Izumchenko, Daria A. Gaykalova, Christine H. Chung, and Elana J. Fertig. Integrated time course omica analysis distinguishes immediate therapeutic response from acquired resistance. Genome Medicine, 10:1–22, 2018.

[51] X. Sun and B. Hu. Mathematical modeling and computational prediction of cancer drug resistance. Brief Bioinform., 19:1382–1399, 2018.

[52] Salvatore Torquato. Random Heterogeneous Materials: Microstructure and Macroscopic Properties. 2002.

[53] J.B. Vermorken, R. Mesia, F. Rivera, E. Remenar, A. Kawecki, S. Rottey, J. Erfan, D. Zabolotnyy, H.R. Kienzer, D. Cupissol, F. Peyrade, M. Benasso, I. Vynnychenko, D. De Raucourt, C. Bokemeyer, A. Schueler, N. Amellal, and R. Hitt. Platinum-based chemotherapy plus cetuximab in head and neck cancer. N Engl J Med., 359:1116–1127, 2008.

[54] F.-G. Wieland, A.L. Hauber, M. Rosenblatt, C. Tonsing, and J. Timmer. On structural and practical identifiability. Current Opinion in Systems Biology, 25:60–69, 2021.

[55] S. Zhang and G. Lin. Robust data-driven discovery of governing physical laws with error bars. Proc. R. Soc. A, 474:20180305, 2018.

